# A systems biology-based molecular model of the protein kinase C life cycle

**DOI:** 10.1101/652628

**Authors:** Naveed Aslam, Farah Alvi

## Abstract

Protein kinase C (PKC) enzymes are a family of kinases that mediate signal transduction originating at the cell surface. Most cell membranes can contain functional PKC enzymes. Aberrations in the PKC life cycle may result cellular damage and dysfunction. For example, some cancerous cells exhibit alterations in PKC activity. Here, we use a systems biology approach to describe a molecular model of the PKC life cycle. Understanding the PKC life cycle is necessary to identify new anticancer drug targets. The PKC life cycle is composed of three key regulatory processes: maturation, activation and termination. These processes precisely control PKC enzyme levels. This model describes the fate of PKC during *de novo* synthesis and PKC’s lipid-mediated activation cycle. We utilize a systems biology approach to show the PKC life cycle is controlled by multiple phosphorylation and dephosphorylation events. PKC processing events can be divided into two types: maturation via processing of newly synthesized enzyme and secondary messenger-dependent activation of dormant, but catalytically-competent enzyme. Newly synthesized PKC enzyme is constitutively processed through three ordered phosphorylations and stored in the cytosol as a stable, signaling-competent inactive and autoinhibited molecule. Upon extracellular stimulation, diacylglycerol (DAG) and calcium ion (Ca^+2^) generated at the membrane bind PKC. PKC then undergoes cytosol-to-membrane translocation and subsequent activation. Our model shows that, once activated, PKC is prone to dephosphorylation and subsequent degradation. This model also describes the role of HSP70 in stabilization and re-phosphorylation of dephosphorylated PKC, replenishing the PKC pool. Our model shows how the PKC pool responds to different intensities of extracellular stimuli? We show that blocking PHLPP dephosphorylation replenishes the PKC pool in a dose-dependent manner. This model provides a comprehensive understanding of PKC life cycle regulation.

## Introduction

Cell membranes are platforms for the transduction of vital signals affecting cellular fate [1–3]. Signal transduction must occur for cells to function normally [1–2]. Protein kinase C (PKC) enzymes comprise a family of enzymes that transduce signals originating from the cell surface [1]. PKC signaling is initiated by receptor-mediated hydrolysis of membrane phospholipids [1–6]. PKC enzymes are regulated at the level of structure, function and subcellular localization [1–9]. PKC enzyme dysfunction has been linked to many human disease pathologies, including cancer, diabetes and heart disease [10–17]. PKC modulates aspects of cellular physiology, such as angiogenesis, cellular motility, proliferation, differentiation and apoptosis [18]. Differential expression and activity of PKC has been linked to most types of cancer, including carcinomas (e.g., breast, colorectal), sarcomas (e.g., glioma), lymphomas (e.g., diffuse large B-cell lymphoma) and leukemias [10–15]. Generally, loss of PKC function has been linked to cancer, whereas enhanced activity has been linked to neurodegeneration [16]. Many groundbreaking investigations during the last two decades have suggested that PKC enzymes have novel therapeutic potential as drug targets in different cancers, heart failure and neurodegenerative diseases [10–19]. Diverse catalytic and regulatory mechanisms exist within the PKC enzyme family, providing a plethora of possibilities for drug target design [18–26].

Cellular function depends on the ability of cells to mount dynamic responses to environmental cues. A critical part of this cellular response is the tight modulation of signaling events [27]. Phosphorylation is a crucial mechanism to transduce environmental signals from the cell membrane to the cytoplasm. Cellular phosphorylation of substrates is controlled by hundreds of kinases and phosphatases. Global cellular functions (i.e. proliferation and apoptosis) and specialized functions (i.e. hormone secretion) maintain homeostasis through phosphorylation and dephosphorylation of signaling substrates. Imbalances in the phosphorylation states of important substrates may result in a disease state, such as cancer or neurodegeneration. Second messengers like Ca^+2^/DAG transmit crucial signaling information that controls cellular fate. PKC family members bind to secondary messengers, which may play a critical role in translating environmental cues into useful and vital intracellular information by determining the phosphorylation status of key downstream substrates. Phosphorylation is key in the regulation of PKC family members because the enzymes are constitutively phosphorylated. PKC enzyme dephosphorylation is agonist-driven [24,25,26]. Novel therapies may be designed to target PKC phosphorylation states.

As PKC dysfunction can result in disease states, it is crucial to understand the underlying molecular mechanisms that control cellular levels of PKC. Understanding downstream signaling of PKC will aid in rational design of new therapeutic strategies. Here, we propose a molecular model of the PKC life cycle based on a systems biology approach. Our model is based on two key observations: (a) phosphorylation controls PKC enzyme stability; and (b) PKC activation and termination are agonist-modulated events [24–26]. According to this model, since PKC is constitutively phosphorylated, PKC dephosphorylation modulates PKC levels and the amplitude of PKC signaling in cells.

PKC protein conformation is tightly controlled. Conformational control is required for PKC to function in a multitude of cell types. PKC conformation may be regulated by two molecular mechanisms: (a) constitutive phosphorylation; and (b) secondary messenger-dependent membrane binding. Newly-synthesized PKC is unstable and is quickly degraded when not modified by phosphorylation. Naïve PKC undergoes a series of ordered phosphorylations in order to avoid degradation [18–20,23,24]. These phosphorylations stabilize PKC, rendering it a catalytically competent and auto-inhibited enzyme [23–24]. Catalytically primed, auto-inhibited PKC is diversely localized within cells. Localization of PKC is maintained by tethering to multimolecular complexes or to other structures [23,24]. Constitutive phosphorylation of PKC serves two main regulatory functions in the life cycle of both conventional and novel PKCs: (a) stabilization of nascent enzyme to prevent degradation; and (b) priming PKC to rapidly activate in response to interaction with secondary messengers [24]. Stabilization of nascent PKC is crucial because the amplitude and duration of signaling downstream of PKC could be linked to the changes in PKC levels. Nascent PKC stabilization may also affect agonist-induced PKC signaling in cells.

This study proposes a comprehensive model of the PKC life cycle from biosynthesis to degradation. Rationale for the model proposed in this study was derived from many of the following experimental observations [1–9,18–21,23–24]: (a) Depleted PKC has been observed in cells lacking kinases, such as PDK1. (b) In addition, nascent PKC phosphorylation at activation site is constitutive. (c) It has been shown that PKC must bind to complexes like mTORC_2_ for phosphorylation to occur at the hydrophobic motifs of PKC. (d) It is known that constitutive phosphorylation regulates PKC stability and catalytic competence. (e) Sustained elevation of the endogenous activator DAG leads to activation, dephosphorylation and down-regulation of PKC. (f) Inhibition of PKC dephosphorylation results in PKC accumulation. (g) PH domain and Leucine rich repeat Protein Phosphatase (PHLPP)-mediated dephosphorylation controls PKC degradation. (h) In the absence of chronic second-messenger cell stimulation, PKC isozyme has a relatively long half-life on the order of days. (i) Finally, a family of 70 kilodalton heat shock proteins called HSP70 proteins are required for the re-phosphorylation of PKC following PKC activation. Re-phosphorylation protects active PKC from degradation, replenishing the cellular pool of PKC for the next cycle.

In this study, we examine the mechanisms by which the cellular PKC pool is established and maintained through constitutive phosphorylation events after *de-novo* synthesis. In addition, we explore possible molecular mechanisms responsible for PKC activation and down-regulation during agonist-induced stimulation. Understanding the PKC life cycle is crucial to designing therapeutics targeting PKC enzymes to alleviate many human diseases. PKC dysfunction has been correlated with a multitude of disease states. For example, PKCβ protein levels are 4-5-fold higher in breast cancer cells than in normal mammary cells [28]. Here, we use a systems biology model of the PKC life cycle to show that PKC down-regulation is dependent on the amplitude and the duration of second messenger stimulation. Our results also suggest that PKC down-regulation depends on dephosphorylation pathways. Specifically, when a cell is exposed to a larger second messenger pulse stimulation, PKC becomes locked in a dephosphorylated, but active, state. In response to this kind of stimulation, almost all the cellular PKC pool is quickly degraded. Degradation occurs because the HSP70-mediated re-phosphorylation process slows under these conditions, and can therefore only minorly contribute to rescue, active and dephosphorylated PKC. Finally, our results demonstrate that blocking the PHLPP-mediated dephosphorylation pathways eliminates agonist-mediated PKC down-regulation, even during highly stimulated cellular conditions.

Our findings indicate that the PKC life cycle is subject to tight temporal modulation that occurs through a complex sequence of phosphorylation events. These molecular events are broadly classified as either constitutive or agonist-induced. Constitutive events are dominant during the early part of the PKC life cycle, when PKC is most unstable. Agonist-induced events are dominant later in the PKC life cycle, during the activation and down-regulation phases. Over the course of the PKC life cycle, naïve PKC transforms into a specific transducer of extracellular information into a multitude of intracellular signals. Using a systems-based approach, we show that the amount of activated PKC affects cellular PKC levels. The relationship between activated PKC levels and the PKC cellular pool may have significant implications for human disease. Our findings also show that heat shock proteins play an important role in the re-phosphorylation and re-stabilization of PKC, even during highly stimulated cellular conditions. Our results indicate that even when cells are highly stimulated, PKC can be rescued from degradation. The amount of re-phosphorylated PKC increases in a dose-dependent manner in response to increasing HSP70 protein expression. The results presented in this study provide a kinetic-mechanistic model for the PKC life cycle. This model may be instrumental in drug design targeting members of the PKC enzyme family.

## Results

### A Model of the PKC Life Cycle

In this study, we propose a model for the PKC enzyme life cycle. This model explains the fate of the nascent PKC molecule. We propose (Figure 1) that temporal regulation of conventional and novel PKCs is based on two phases: 1. constitutive phase; 2. agonist-induced activation/down-regulation/termination phase. In this model, PKC could be either an mRNA (PKC mRNA) or a protein (PKC). The PKC protein can be in one of six states: nascent enzyme at membrane (PKC), enzyme phosphorylated at activation site through PDK1-mediated molecular event at membrane (PKC.P_A_), the complex of PKC.P_A_ with mTORC_2_ at the membrane (C.PKC.P_A_), the enzyme-complex phosphorylated at activation, hydrophobic and turn motifs and located in the cytosol (C.PKC.P_A_.P_H_.P_T_); the activated & phosphorylated enzyme at the membrane (PKC.P_A_.P_H_.P_T_)^A^; and the dephosphorylated but active molecule located in the cytosol (PKC^A^). According to this model, brief protein synthesis stimulation leads to generation of naïve PKC. Newly synthesized, unphosphorylated PKC is highly unstable and only loosely tethered at the cell membrane. Naive PKC is an exposed pseudo-substrate with an accessible C-terminal tail. According to the model, constitutively-active PDK_1_ docks with the C-terminal of nascent PKC and modulates PKC phosphorylation at activation loop (P_A_). Constitutive phosphorylation stabilizes the previously-labile naïve enzyme and triggers binding to the mTORC2 complex at the cellular membrane. This binding event, in turn, catalyzes a series of sequential autophosphorylations on PKC’s C-terminal tail, hydrophobic motif and turn motif. These phosphorylations are denoted as P_H_ and P_T_. These three constitutive ordered phosphorylations generate a mature, phosphatase/protease resistant, but catalytically competent molecule denoted as C.PKC.P_A_.P_H_.P_T_. This catalytically-competent molecule is then localized in the cytosol and stored there. The C.PKC.P_A_.P_H_.P_T_ form of PKC is stable and inactive, but catalytically competent and functionally responsive to second messenger changes inside the cell or at the membrane compartment. According to this model, the first phase of the PKC life cycle is constitutive phosphorylations. During this phase, after a brief protein synthesis pulse, the naïve molecule is converted into a stable and inactive species, but is still responsive to second-messengers. In the first phase of the life cycle, PKC is stored at many locations throughout the cytosol. Our model emphasizes how PKC functionality and signaling depends on constitutive processing events, leading to its stabilization and catalytic competence?

**Figure 1:**
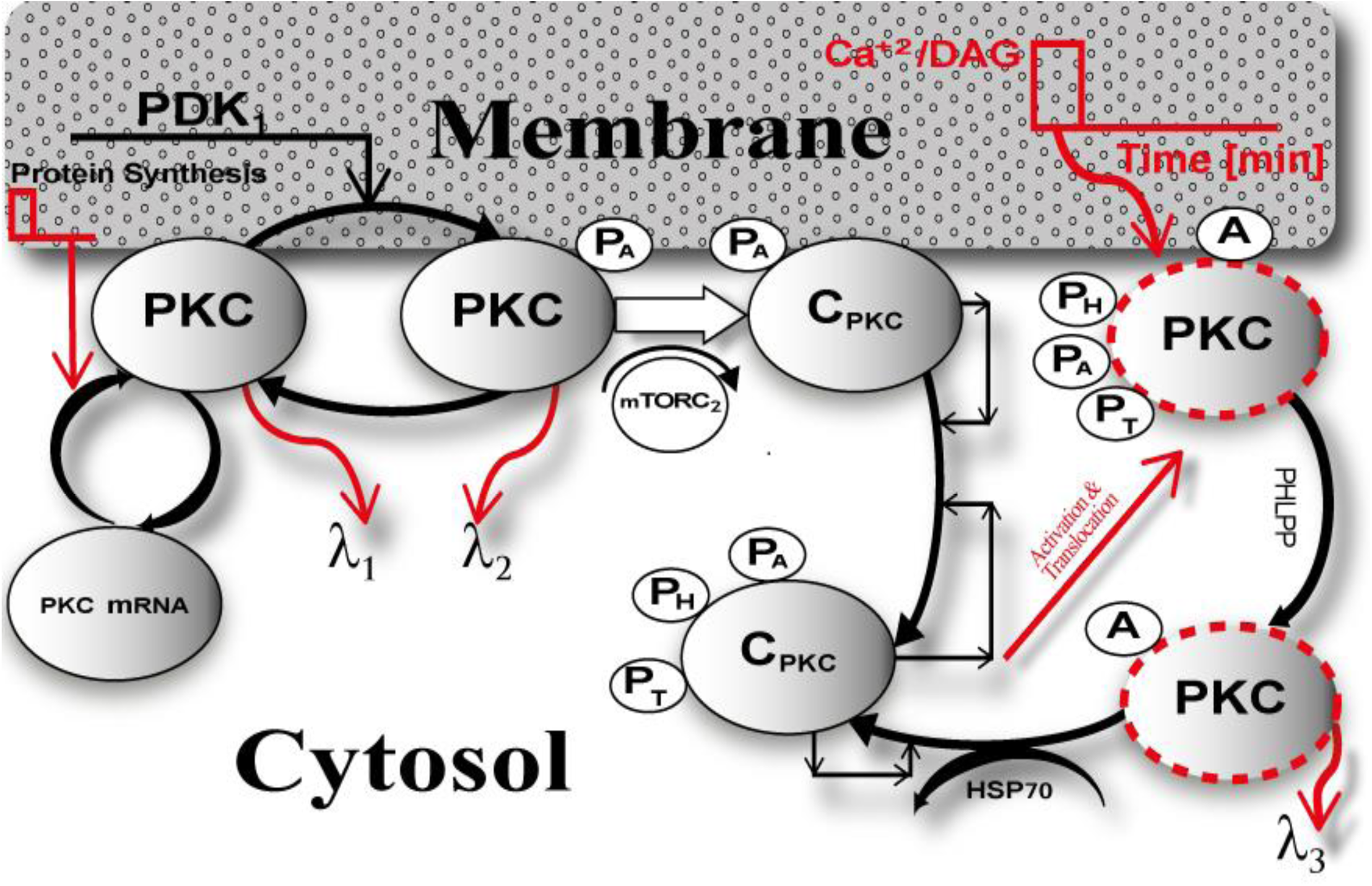
The life cycle model of PKC enzyme. This model explains the fate of naïve, PKC molecule. According to this model the PKC life cycle has two phases a) constitutive phase and b) agonist-evoked phase. The constitutive phase of PKC life cycle is involved in stabilization and maturation processing of newly synthesized enzyme. According to this model the newly synthesized unphosphorylated cPKC isozymes are highly unstable and are only loosely tethered at the membrane with exposed pseudo substrate and accessible C-terminal tail. This model describes that constitutively active upstream kinase, PDK1, docks onto the C-terminal tail of this newly synthesized PKC molecule, thus allowing efficient phosphorylation of activation loop (P_A_). This initial phosphorylation stabilizes the labile enzyme and triggers two sequential phosphorylation events on the C-terminal tail i.e., hydrophobic & turn motif phosphorylations (P_H_ & P_T_). These three constitutive ordered phosphorylations generate a mature, phosphatase/protease resistant molecule (CPKC.P_A_.P_H_.P_T_). This fully phosphorylated mature molecule then localizes to cytosol, where it is maintained in an inactive and phosphatase resistant conformation. This model thus suggests that after a brief protein synthesis pulse newly synthesized PKC molecule is stabilized and stored in cytosol through a sequence of constitutive events. This model also points out that PKC function is pretty much dependent-on the phosphorylations at activation, hydrophobic and turn motifs. The agonist-evoked phase of signaling involves second messenger Ca^+2^/DAG modulated activation and subsequent termination of PKC signaling through dephosphorylation and degradation. According to this model a brief second messenger pulse at the membrane compartment can lead to translocation and activation of mature PKC enzyme and second messenger binding at membrane transforms the inactive mature molecule to active specie i.e., C.PKC.P_A_.P_H_.P_T_ ^A^. This active molecule is prone to dephosphorylation and upon PHLPP-mediated dephosphorylation event relocates back to cytosol where either it can be either degraded or re-phosphorylated through hsp70 and autophosphorylation-mediated events. This model suggests that chronic activation of cPKC molecule could possibly result in the complete down-regulation of enzyme. This model describes that PKC signaling lifetime is composed of synthesis, maturation, activation and termination events.

This model suggests that, during the second phase of the life cycle, inactive, phosphatase/proteasome-resistant PKC species are activated through second messengers Ca^+2^ and DAG. A brief second-messenger pulse at the membrane generates second messengers Ca^+2^/DAG. This pulse, in turn, leads to translocation, activation, down-regulation and subsequent termination of PKC. The model shows that a brief second-messenger pulse of DAG in the membrane compartment leads to the translocation of mature PKC from the cytosol to the membrane. At the membrane, binding with a second-messenger converts inactive PKC to active, but phosphatase-sensitive PKC. This form of PKC is denoted as (PKC.P_A_.P_H_.P_T_)^A^. In this stage, PKC is active but prone to dephosphorylation. PHLPP-mediated dephosphorylation then relocates PKC back to the cytosol. Active but dephosphorylated PKC^A^ could also be targeted for degradation or re-phosphorylation by HSP70-mediated rescue and autophosphorylations. Our model suggests that chronic activation of PKC may result in complete dephosphorylation and down-regulation of PKC. PKC’s life cycle involves *de-novo* synthesis, maturation, activation and termination. Each of these events affects PKC levels and, thus, the agonist-mediated amplitude and duration of active PKC signaling.

### PKC life cycle: role of protein synthesis and second messenger modulated activation

Using our model, we determined the PKC signaling characteristics like PKC levels and the duration of non-negligible concentrations during protein synthesis, second messenger-modulated activation, and down-regulation. Signaling characteristics were determined using the levels of total PKC and the relative distribution of all six enzyme species during different phases of the PKC life cycle. We also measured how long total PKC levels and the levels of different forms of PKC are non-negligible? Here, dashed red lines indicate stimulated conditions like protein synthesis and second messenger-mediated activation. Blue solid lines indicate conditions where there was no stimulation. In these simulations, protein synthesis is initiated by a 10-minute pulse, leading to the generation of naïve, unstable PKC. Second messenger stimulation, or a pulse mimicking DAG generation, is applied 50 minutes into each simulation. The second messenger pulse is applied for 15 minutes. Four different levels of DAG intensity are used in the pulse stimulation. The DAG intensity is set at 0.0005 (nM) and is linked to dashed line representing: a-----a, intensity set at 0.005 (nM) and is linked to b----b, intensity set at 0.05 (nM) and is linked to c----c and DAG intensity set at 0.5 (nM) and is linked to d-----d. These different strengths of second messenger pulse indicate different levels of PKC activation (Figure 2: **dashed red lines, a----a, b----b, c----c and d----d**). In the absence of a protein synthesis pulse, there is no *de-novo* PKC synthesis and the system is fixed in the basal state (Figure 2: **solid blue line**). Additionally, in the absence of a second messenger DAG pulse or low intensity stimulation pulse, there is very little change in total PKC levels (Figure 2: **red dashed line a-----a)** and transformations into activation states of [PKC.P_A_.P_H_.P_T_]^A^ and PKC^A^ are negligible.

**Figure 2:**
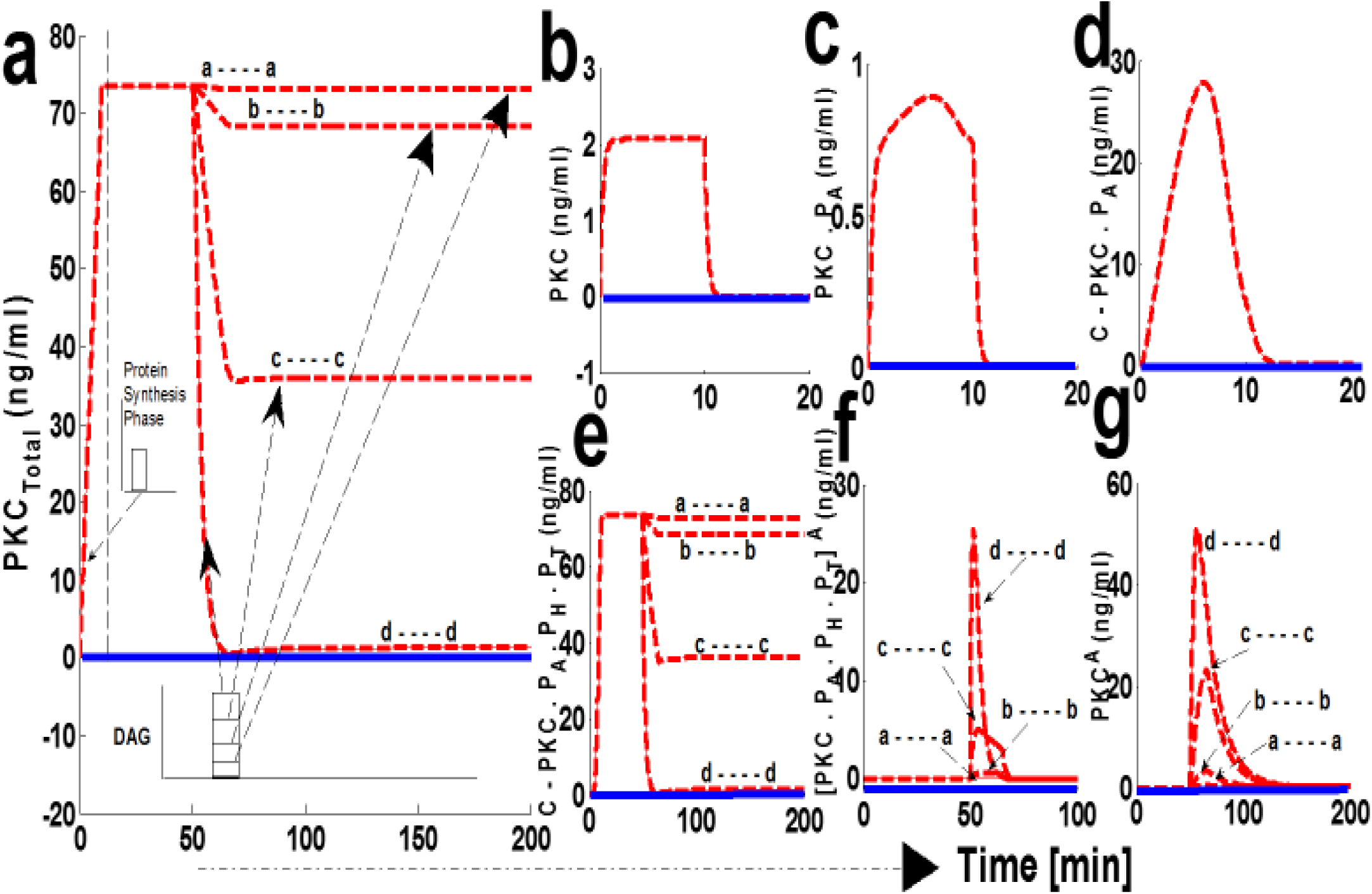
The numerical simulations illustrating the role of protein synthesis and second messenger-mediated activation on cPKC life cycle. These results indicate that initial application of a 10-minute protein synthesis pulse leads to the generation of a naïve, and unstable enzyme which is quickly stabilized through constitutive phosphorylation events at the activation, turn and hydrophobic sites. Once stable and phosphatase/proteasome resistant this inactive form of cPKC is stored in cytosol. Here, dashed red lines indicate stimulation (**protein synthesis as well as second messenger**) and blue solid line indicate non-stimulated conditions. (a) Total PKC enzyme during protein synthesis and second messenger mediated stimulation. These results show that application of a brief pulse (**dashed-line region**) shows the quick generation of cPKC enzyme and stabilization. These results also show that at later time point (**i.e., 50 minutes in the simulations**) the application of a second messenger pulse (**15-min pulse mimicking Ca^+2^/DAG**) leads to the activation, termination and down-regulation of cPKC enzyme. These results high light that degree of cPKC down-regulation is influenced by the duration and strength of second messenger pulse. Here, four different stimulation strengths of pulse are used i.e., levels of second messenger DAG are set at 0.0005 nM (**a------a**), DAG set at 0.005 nM (**b-----b**), DAG set at 0.05 nM (**c----c**) and DAG set at 0.5 nM (**d-----d**) (**from lowest to highest strength levels**). These results elucidate that at lowest second messenger stimulation strength i.e., 0.0005 nM there is no down-regulation of cPKC and at highest stimulation strength i.e., 0.5 nM cPKC is completely down-regulated. (b) concentration of naïve, newly synthesized unstable form of cPKC. (c) concentration of PKC species phosphorylated at activation site. (d) concentration of PKC complex C.PKCP_A_. (e) concentration of PKC complex, phosphorylated at all three sites i.e., activation, turn and hydrophobic sites. These results show that complex C.PKC.P_A_.P_H_.P_T_ is down-regulated by higher stimulation levels of second-messenger DAG. (f) concentration of active complex PKC.P_A_.P_H_.P_T_.^A^ after second-messenger binding. Again, these results indicate that activation levels of this complex are dependent on the levels of second-messenger stimulation. (g) concentration of dephosphorylated but active PKC^A^ molecule.

When a protein synthesis pulse is applied the total amount of PKC quickly increases to 73 ng/ml, then stabilizes (Figure 2a). This result shows that, after de-novo synthesis, newly synthesized PKC is quickly processed through a constitutive mechanism, i.e., three ordered phosphorylations (Fig. 2b, 2c and 2d). PKC is then stored in the cytosol as a stable enzyme competent for signaling, denoted in our model as C.PKC.P_A_.P_H_.P_T_ (Figure 2c). This first phase of the PKC life cycle lasts between 0-50 minutes (till the application of second messenger pulse stimulation) and illustrates the role of PDK_1_, mTORC_2_, and autophosphorylations in converting labile PKC to a stable, mature, and inactive but catalytically-competent molecule. Stable PKC is stored in the cytosol where it then relays second messenger information from the cell surface to key intracellular targets on the interior of the cell. During this first phase of the PKC life cycle the concentration of both the active species i.e., [PKC.P_A_.P_H_.P_T_]^A^ and PKC^A^ are negligible, as no second messenger stimulation has been applied. Both these species remain at basal level concentration during this phase.

These numerical simulations also illustrate that at a later time point, i.e., 50 minutes into simulation, when a second messenger DAG pulse is introduced, varying levels of the pulse leads to changes in total PKC levels (Figure 2a **dashed red lines a----a, b----b, c----c**). Levels of three states of cPKC isozymes i.e., C.PKC.P_A_.P_H_.P_T_ (Figure 2c), PKC.P_A_.P_H_.P_T_.^A^ (Figure 2f) and PKC^A^ (Figure 2g) also change. These results show that second messenger-modulated stimulation initiates the second phase of cPKC life cycle. This phase could possibly involve translocation, activation, termination and down-regulation events. These results highlight how cPKC down-regulation depends on the duration and strength of the second messenger pulse? In our simulation, we used four different levels of DAG stimulation. Our results show that at the lowest level of second-messenger stimulation, 0.0005 (nM), there is no down-regulation of cPKC. At the highest level of stimulation, 0.5 (nM) cPKC is completely down-regulated (Figure 2a; a---a; b---b; c----c; & d----d). Interestingly, these results also show that PKC down-regulation depends upon PKC.P_A_.P_H_.P_T_.^A^ and PKC^A^ states (Figure 2f **and** 2g). In the case of a strong second-messenger pulse (**d-----d**), the concentration of PKC.P_A_.P_H_.P_T_.^A^ and PKC^A^ reach their highest levels, leading to significant degradation of PKC. These results also underscore the importance of balance between PKC re-phosphorylation and degradation in controlling the overall PKC concentration after second messenger stimulation.

As part of our investigation we also analyzed how second messenger pulse duration affects the PKC life cycle. We set DAG pulse intensity to 0.0005(nM) and varied the duration of this low-level pulse. In these simulations (Figure 3) we used five different durations of pulse stimulation: 15 min. (a------a), 100 min. (b-----b), 150 min. (c----c), 200 min. (d-----d) and 316.66 minutes (e-----e). Our results show that increasing the duration of the second messenger pulse has only a moderate effect on the PKC life cycle (Figure 3). For lower intensity stimulation, the degree of down-regulation is almost negligible when pulse stimulation is set at 15 minutes (Figure 3 a-----a) and is only moderate when pulse stimulation is set at 316.66 minutes (Figure 3 d-----d).

**Figure 3:**
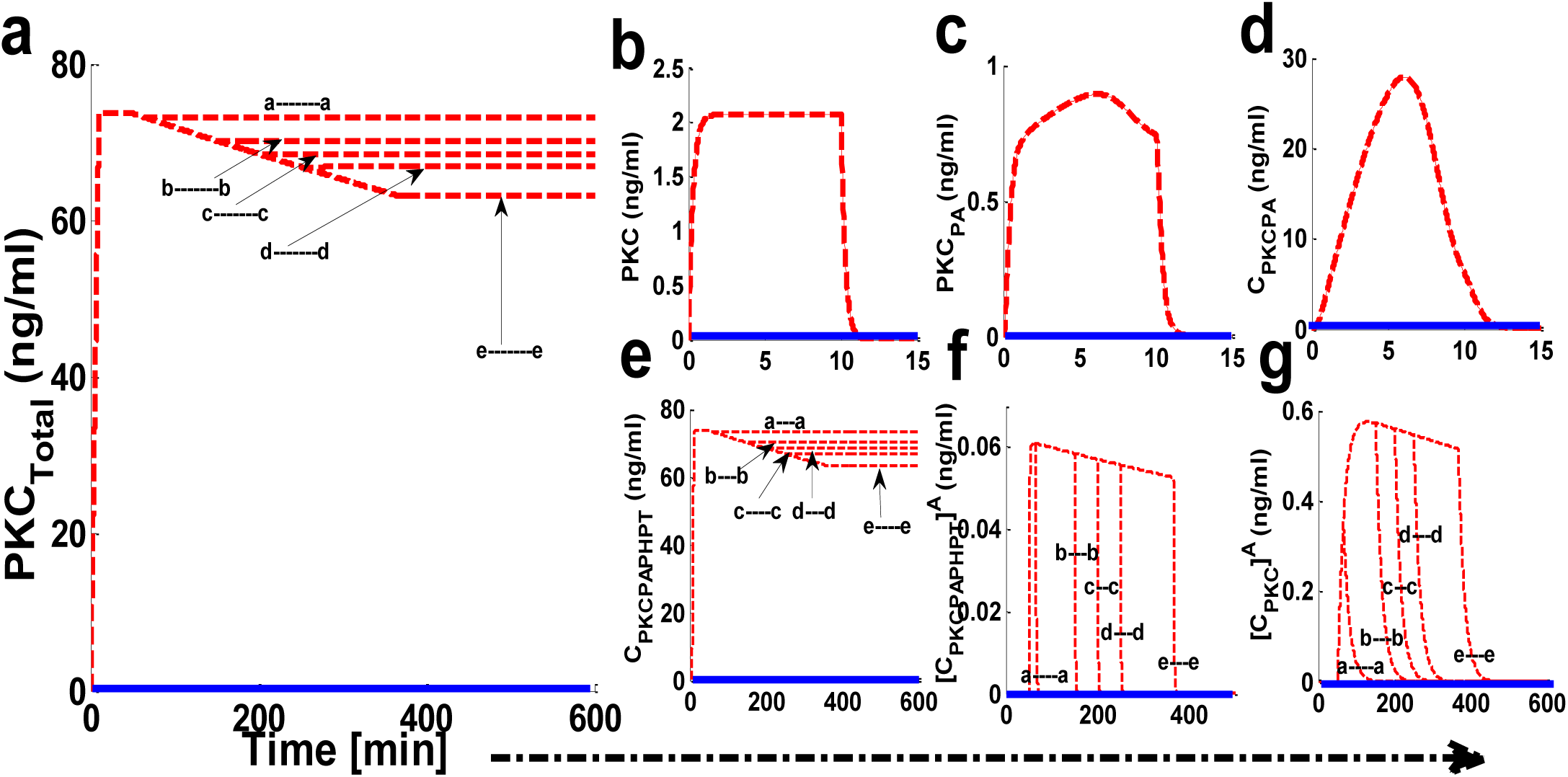
The effect of second messenger-modulated activation duration on the life cycle characteristics of PKC. These results indicate that initial application of a 10-minute protein synthesis pulse leads to the generation of a naïve, and unstable enzyme which is quickly stabilized through constitutive phosphorylation events at the activation, turn and hydrophobic sites. Once stable and phosphatase/proteasome resistant this inactive form of cPKC is stored in cytosol. Here, solid red lines indicate stimulation (**protein synthesis as well as second messenger**) and blue solid line indicate non-stimulated condition. These results show that during lower intensity second messenger stimulation increasing the duration of pulse can influence the life cycle characteristics of enzyme. However, the degree of this influence is not significant. (a) Total PKC enzyme during protein synthesis and second messenger mediated stimulation. These results show that application of a brief pulse protein synthesis pulse (10 minutes) shows the quick generation of cPKC enzyme and stabilization. These results also show that at later time point (i.e., 50 minutes) the application of a lower intensity second messenger pulse (15-min pulse mimicking Ca^+2^/DAG generation; DAG = 0.0005 nM) leads to the activation, termination, down-regulation and restoration of cPKC enzyme. These results high-light that cPKC signaling characteristics are only moderately influenced by the duration of second messenger pulse. Here, five different durations of pulse stimulation are used i.e., second messenger DAG pulse stimulation set at 15 min. (**a------a**), 100 min. (**b-----b**), 150 min. (**c----c**), 200 min. (**d-----d**) and 316.66 minutes (**e-----e**) (from shorter to longer duration periods). These results elucidate that at shorter pulse period i.e., 15 minutes degree of down-regulation is only minimum and at longest pulse period i.e., 316.66 minutes the degree of down-regulation is only moderate. (b) The concentration of naïve, newly synthesized unstable form of cPKC. (c) The concentration of PKC species phosphorylated at activation site. (d) The concentration of PKC complex C.PKC.P_A_. (e)The concentration of PKC complex, phosphorylated at all three sites i.e., activation, turn and hydrophobic sites. (f) The concentration of active complex PKC.P_A_.P_H_.P_T_^A^ after second messenger binding. (g) Concentration of dephosphorylated but active PKC^A^ molecule.

### Effect of blocking PKC dephosphorylation

We next determined whether blocking PKC dephosphorylation could influence the PKC life cycle. In these simulations, the protein synthesis protocol is the same as described in the previous section. We applied three sequential pulses of DAG during simulations (time = 4000 s; =12,000 s; & = 20,000 s). The strength of these sequential pulses was 0.5 (nM) with a fixed duration of 15 minutes. During these activation pulses dephosphorylation is inhibited by completely blocking the parameter k_17_. Our results show that blocking dephosphorylation prevents down-regulation of cPKC even at higher stimulation levels (Figure 4 **dashed red lines for total enzyme and inset for different states of the enzyme**). These results show that, if dephosphorylation is blocked, PKC levels are maintained. This shows that high-intensity, activation-induced down-regulation of PKC can take place through dephosphorylation. These results imply that if dephosphorylation could be controlled, PKC down-regulation could possibly be fine-tuned. Interestingly, sequential application of second messenger pulses can lead to decreases in C.PKC.P_A_.P_H_.P_T_ concentration and increases in PKC.P_A_.P_H_.P_T_ ^A^ concentration (Figure 4 **inset dashed red lines for different states of the enzyme**). However, once DAG is removed, C.PKC.P_A_.P_H_.P_T_ and total PKC levels are restored back to their original values. These results emphasize the essential role of different enzyme states in response to second messenger stimulation.

**Figure 4:**
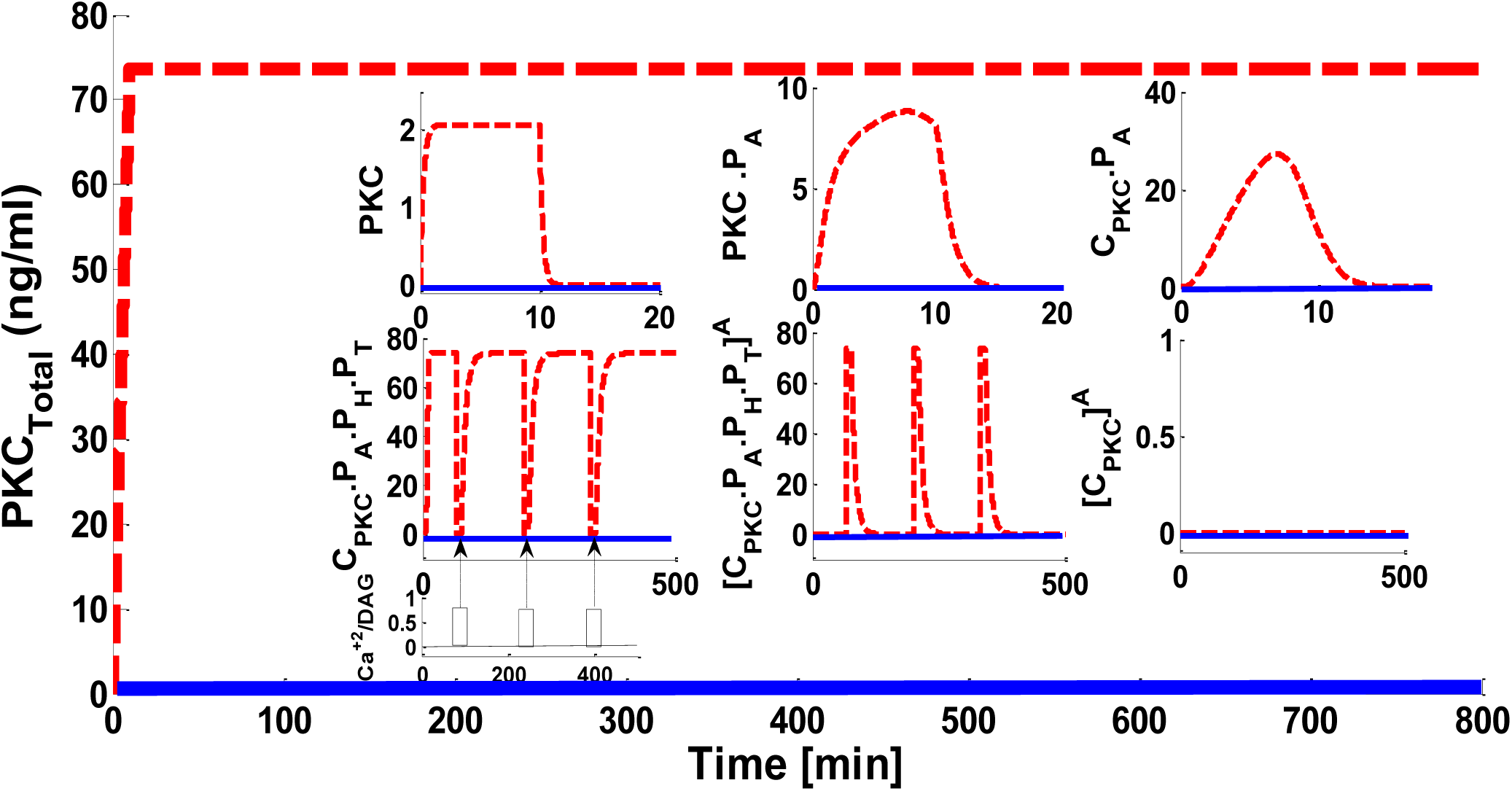
The effect of 100% blocking of dephosphorylation rate i.e., “k_17_”, on the cPKC life cycle. These results show that 100% blocking of dephosphorylation rate prevents the down-regulation of total PKC, even at higher stimulation levels. Here, application of initial protein synthesis stimulation leads to the quick generation and constitutive stabilization of cPKC enzymes. After initial protein synthesis stimulation three identical pulse stimulations of second messenger DAG are applied at time points t = 4000 secs; t = 12,000 secs; and t = 20,000 secs. Each of these pulse stimulations is applied for 15 minutes and during their application the dephosphorylation rate k_17_ was completely blocked. These results indicate that blocking the dephosphorylation did not result in the down-regulation of total PKC enzyme as well as the CPKC.P_A_.P_H_.P_T_. The inset shows that application of a second messenger stimulation results in decrease of CPKC.P_A_.P_H_.P_T_ and increase in PKC^A^ but after second messenger is removed the CPKC.P_A_.P_H_.P_T_ levels are restored indicating that down-regulation is very much dependent on the dephosphorylation pathways.

### Effects of sequential intermediate strength second messenger pulse stimulations on the PKC life cycle

As shown in previous sections, PKC down-regulation can be controlled through varying the strength and duration of second messenger pulse stimulation. Next, we set out to investigate whether sequential intermediate strength second messenger pulses affect the PKC life cycle. Three identical strength second messenger pulse stimulations were applied at three different time points. The first pulse was applied at 50 minutes; the second pulse was applied at 220 minutes; and the third pulse was applied at 360 minutes. For all three pulses, DAG pulse strength is set at an intermediate level (Fig 2a. inset pulse b---b; DAG = 0.005 au). Our results indicate that second messenger-induced destabilization and down-regulation takes place when pulse stimulation is applied. PKC down-regulation occurs in response to pulse stimulation, but overall down-regulation depends on the number of pulses applied (Figure 5a). These results also indicate three downregulation events occur in response to three pulse stimulations (Figure 5). Interestingly, the extent of down-regulation decreases with each sequential pulse event. After the first pulse, PKC levels drop almost 10 ng/ml. In comparison, for the third pulse, PKC levels drop is only 5 ng/ml. This observation shows that, as the number of pulses increases, the net effect of the pulses on PKC degradation and down-regulation also decreases. This occurs even though the strength of each pulse remains constant. The same phenomenon was observed for PKC.P_A_.P_H_.P_T_ ^A^ and PKC^A^ (Figure 5f **and** 5g).

**Figure 5:**
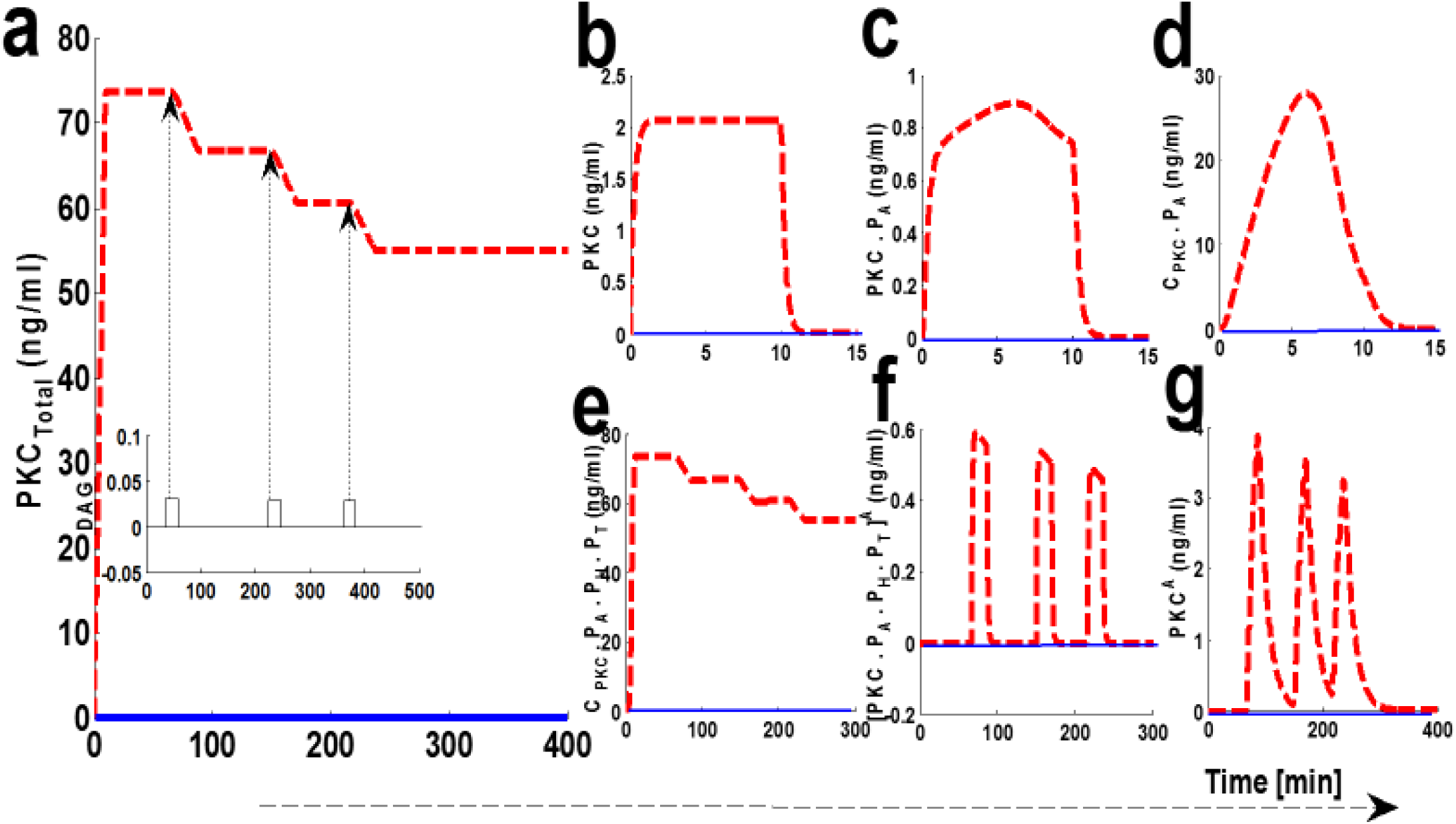
The effect of sequential application of second messenger stimulation on the down-regulation characteristics of the cPKCs. Here, dashed red lines represents the stimulation whereas, the solid blue line represents non-stimulated conditions. Here, three equal pulses of second messenger are applied in a sequential manner. The pulse 1 is applied at t = 50 minutes; the pulse 2 is applied at t = 220 minutes; and the pulse 3 is applied at t = 360 minutes. For all these three pulses second messenger levels are set at DAG = 0.005 nM; These results show the second messenger-induced destabilization and down-regulation of cPKCs. These results also show that with application of each pulse, a certain amount of enzyme is down-regulated and total PKC levels are stabilized at new levels. These results indicate that during simulations three down-regulation events are observed each in response to a second messenger pulse at t = 50 minutes; t = 220 minutes; & t = 360 minutes. (a) Total PKC enzyme during protein synthesis and second messenger mediated stimulation. (b) The concentration of naïve, newly synthesized unstable form of cPKC. (c) The concentration of PKC species phosphorylated at activation site. (d) The concentration of PKC complex C.PKC.P_A_. (e)The concentration of PKC complex, phosphorylated at all three sites i.e., activation, turn and hydrophobic sites. (f) The concentration of active complex PKC.P_A_.P_H_.P_T_^A^ after second messenger binding. (g) Concentration of dephosphorylated but active PKC^A^ molecule.

### How HSP70 proteins may influence the PKC life cycle?

Previous studies indicate heat shock proteins may prevent agonist-associated dephosphorylation and subsequent down-regulation of cPKC enzymes [42]. PKC-HSP70 interactions could be important in certain cancers, where overexpression of HSP70 may prevent the loss-of-function (LOF). LOF is thought to be associated with the down-regulation of cPKC enzymes. We set out to investigate whether overexpression of heat shock proteins could rescue PKC from degradation. In order to study this question, we developed five simulations: wild-type with no-hsp overexpression; 10x overexpression; 40x overexpression; 100x overexpression, and 200x overexpression (Figure 6). In these simulations, second messenger pulse strength is set to 0.5 (nM). Here, dashed red lines show the stimulated conditions, whereas the solid blue lines indicate non-stimulation conditions. These results indicate that when there is no hsp70 overexpression, second messenger stimulation can lead to complete down-regulation of PKC (Figure 6-a**, dashed red line, no hsp70 overexpression**). Interestingly, when simulations are carried out with varying levels of hsp70 overexpression, the degree of cPKC enzyme down-regulation can be reversed in a dose-dependent manner (Figure 6a). These results indicate the possible role of hsp70 expression in PKC rescue. These results show that heat shock protein expression may be instrumental in PKC recovery. C.PKC.P_A_.P_H_.P_T_, also shows complete down-regulation under no hsp70 overexpression and subsequent reversal in a dose-dependent manner upon hsp overexpression (Figure 6b). Interestingly, the overexpression of hsp70 reduces the activation & duration of both PKC.P_A_.P_H_.P_T_ ^A^ and PKC^A^ in a dose-dependent manner, (Fig 6c and d). Heat shock proteins rescue enzymes from degradation and induce re-phosphorylation and autophosphorylation. These results confirm heat shock proteins could be the key to maintaining the cellular pool cPKC after activation. Heat shock proteins can specifically interact with active and dephosphorylated cPKC enzymes, thus reducing cPKC concentration.

**Figure 6:**
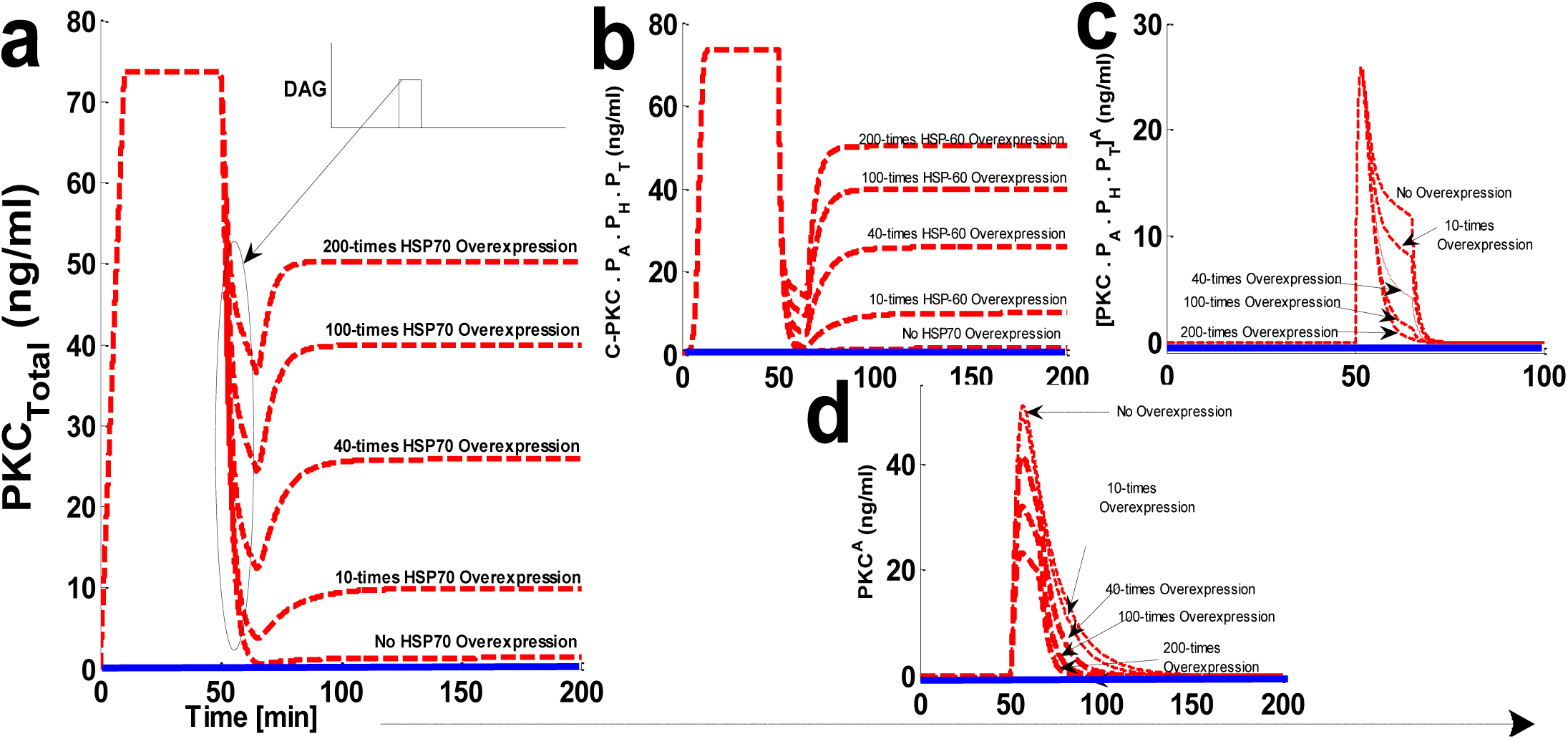
The effect of hsp70 overexpression on the cPKC life cycle. Here, the dashed red line shows stimulated conditions whereas, solid blue line show non-stimulated conditions. (a) The concentration of total PKC enzyme. Here, under the no hsp70 overexpression condition the strong second messenger stimulation leads to the complete down-regulation of cPKC enzyme. Interestingly, when these simulations are run with hsp70 overexpression the degree of down-regulation after second messenger stimulation is reduced in a dose-dependent manner. These results indicate the possible role of hsp70 expression in re-stabilization of active but dephosphorylated PKC enzyme. (b) The concentration of C-PKCP_A_.P_H_.P_T_ with no hsp70 overexpression and dose-dependent reversal of C.PKC.P_A_.P_H_.P_T_ concentration with hsp70 overexpression. (c) The concentration of [PKC.PA.PH.PT]^A^ with no hsp70 overexpression and dose-dependent reduction of [PKC.PA.PH.PT]^A^, concentration with hsp70 overexpression. (d) The concentration of PKC^A^, with no overexpression and dose-dependent reduction in concentration and time persistence with hsp70 overexpression.

### Effects of blocking PHLPP on the cPKC life cycle

The protein phosphatases PHLPP_1_ and PHLPP_2_ are endogenous negative regulators of cPKC signaling. Both these phosphatases play a key role in regulating the dephosphorylation process of cPKC in different cell types. Studies indicate that both phosphatases can bind and dephosphorylate PKCβII on the hydrophobic motif, thus shunting it towards degradation. These phosphatases have also been linked to cancer. We investigated how blocking PHLPP signaling affected the cPKC life cycle? We addressed this question by developing five simulations mimicking the cPKC life cycle. Our simulations include no PHLPP blocking, 50% blocking, 90% blocking, 95% blocking, and 100% blocking. Here, the dashed red line shows activation upon second messenger pulse stimulation. The solid blue line represents no stimulation. DAG levels were set at 0.5 (nM). With no PHLPP blocking cPKC is completely down-regulated (Figure 7a **dashed redline, No PHLPP blocking**). This shows cPKC down-regulation can be reduced in a dose-dependent manner culminating when100% PHLPP blocking results in no down-regulation. Even with high intensity stimulation, the complete blocking of PHLPP results in no down-regulation (Fig 7a). C.PKC.P_A_.P_H_.P_T_ levels increase as PHLPP blocking increases. 100% PHLPP blocking increases the concentration and duration of C.PKC.P_A_.P_H_.P_T_^A^. No PHLPP blocking leads to smaller increases in C.PKC.P_A_.P_H_.P_T_^A^ concentration and a shorter duration of persistence (Figure 7c).

**Figure 7:**
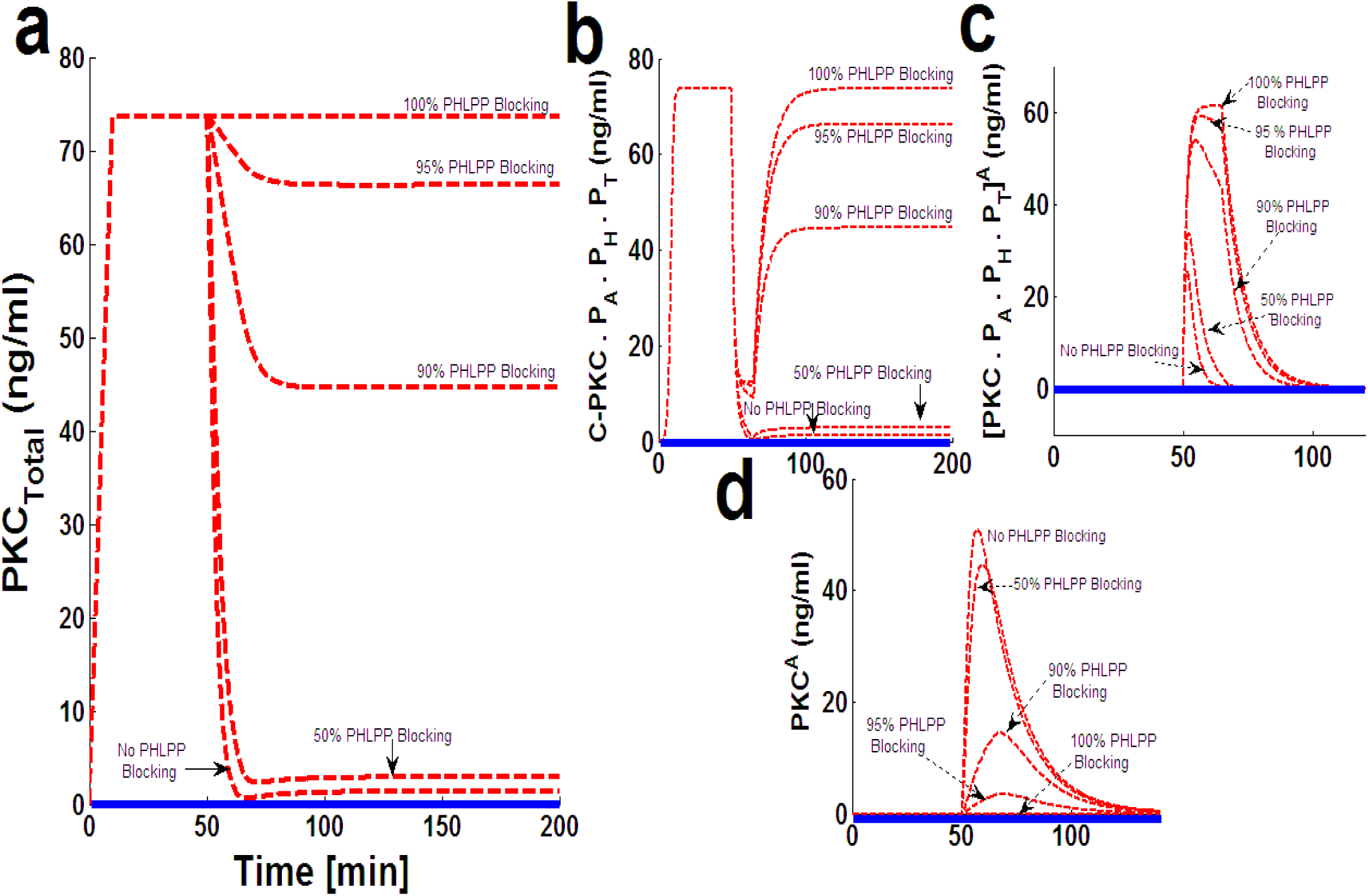
The effect of PHLPP blocking on the down-regulation of cPKC enzyme after second messenger-induced activation. Here, dashed red lines show the stimulation whereas, the solid blue lines show the non-stimulation conditions. (a) The concentration of total PKC. These results indicate that after second messenger stimulation (second messenger DAG = 0.5 nM, with pulse duration of 15 minutes), the cPKC is completely down-regulated when there is no blocking of PHLPP. These results show that degree of cPKC destabilization and its down-regulation is reduced in a dose-dependent manner with increasing the blocking levels of PHLPP. These results indicate that with 100% blocking of PHLPP there is no down-regulation of cPKC, even with high strength second messenger pulse stimulation. (b) concentration of C.PKC.P_A_.P_H_.P_T_ with no blocking and different blocking levels of PHLPP. (c) Concentration of PKCP_A_.P_H_.P_T_.^A^ with no PHLPP blocking and dose-dependent blocking (d) Concentration of PKC^A^ with no PHLPP blocking and different levels of PHLPP blocking.

## Discussion

Over the past two decades, numerous metabolic and signaling pathways have been linked to enzymes of the PKC family [31–41]. PKC family members are critical for assembly of key transduction complexes. These complexes translate environmental cues into critical physiological processes regulating apoptosis, proliferation, gene expression, cell migration, differentiation and cell survival. Due to the PKC enzymes’ wide range of influence on physiological processes, members of this enzyme family could serve as significant targets for drug development. They may lead to novel therapies for human diseases such as cancer, heart failure and neurodegenerative conditions. The functional role of cPKCs in cancer is not understood. In order to better understand the complex functional behavior of PKC in cancer, it is important to understand the regulatory processes involved in PKC modulation. Dissecting the molecular underpinnings of relevant regulatory events linked to individual PKC isoforms is key. Understanding the variations of these regulatory processes across the cell types and even between the tumors could be key in establishing safe and effective therapeutic strategies targeting PKC family enzymes.

Our study focused on understanding the PKC life cycle through a systems biology approach. Here, we proposed a computational model based on elementary kinetics to describe different phases of the PKC life cycle. The proposed model accounts for the maturation, activation and termination phases of the enzyme life cycle. We investigated a general hypothesis that levels of PKC enzyme, and hence the duration and amplitude of associated signaling in cells, are precisely controlled by three key regulatory phases. Any dysfunction in PKC regulation could possibly be linked to disease conditions like cancer. Our proposed life cycle model is based on observations noting that newly-synthesized PKC is unstable, and subsequently processed through constitutive ordered phosphorylations. Activation and termination are agonist-modulated processes and depend on the strength and duration of agonist stimulation.

The proposed network model of PKC life cycle suggests that duration and levels of PKC signaling are regulated by key enzyme interactions during stabilization, activation and down-regulation phases. According to this model, the stabilization and maturation of newly synthesized molecule is modulated by constitutively active PDK1, mTORC_2_ complex and autophosphorylations. Similarly, the translocation of PKC from the cytosol to the cell membrane and second messenger Ca^+2^ and DAG binding play key roles in activation. The down-regulation phase is modulated by phosphatases PHLPP1 and PHLPP2. Interestingly, re-phosphorylation is modulated by hsp70, which can rescue PKC from degradation and target it for storage in an auto-inhibited form in the cytosol. Our results suggest that the duration and strength of cPKC signaling in a multitude of cell types may be modulated by PDK1, mTORC2, PHLPP1 and hsp70. Our results are consistent with experimental observations showing the effects of these kinases and phosphatases on PKC signaling during different phases of the PKC life cycle.

Our study shows that PKC signaling can be fine-tuned by the duration and amplitude of agonist-induced stimulation (Figure 2 **and** 3). Our results show that for small to moderate activation stimuli the pool of total PKC remains unchanged or changes are relatively small (Figure 2a: a----a & b----b). However, high intensity stimuli can lead to complete down-regulation of PKC through activation and dephosphorylation (Figure 2a, e, f & g: c----c & d----d). This occurs because, during the application of higher intensity stimuli, a larger pool of enzyme is activated (Figure 2f: c-----c & d-----d). A larger pool of active enzyme results in much higher concentrations of active, but dephosphorylated, enzyme (Figure 2g: c-----c & d-----d). These results link the larger pool of active and dephosphorylated PKC to higher degrees of down-regulation. Our model predicts that in the wake of high intensity stimuli, endogenous heat shock protein-induced restoration and recovery of cPKC has only limited influence. When there is a small pool of active and dephosphorylated enzymes, the endogenous heat shock-mediated restoration may completely or partially recover PKC from degradation. However, for much larger pools, the enzyme degrades more quickly then it can be re-phosphorylated and replenished in the cytosol by endogenous recovery and restoration factors. This model suggests a delicate balance between degradation and re-phosphorylation pathways controls the degree of PKC down-regulation after agonist-mediated activation.

Our results also analyze the effects of sequential stimulations and complete blocking of dephosphorylation pathways on PKC activation and down-regulation. Dephosphorylation events are critical for targeting active molecules towards degradation. However, down-regulation after high intensity second messenger stimulation might be completely avoided if dephosphorylation pathways are blocked. Our results show that enzyme down-regulation could be completely avoided when dephosphorylation is blocked even under high intensity, sequential second messenger-mediated activation stimuli (Figure 4). Our results also predict that, when dephosphorylation pathways are not blocked, the sequential application of high-intensity stimuli can lead to successive enzyme down-regulation events (Figure 5). However, the degree of down-regulation is reduced with the application of each pulse in a series, despite the strength and duration of the pulses remaining identical (Figure 5, a, e, f, g). This raises an interesting possibility that, initially when PKC levels are high, it might be easier to down-regulate the enzyme. With each cycle the degree of down-regulation could be reduced, raising the possibility that PKC might persist for longer periods after certain initial stimulations in the absence of new protein synthesis or PKC transcription.

In our model, we are not attempting to simulate PKC protein synthesis in a detailed manner, we only intend to mimic the complex signaling cascade involved in new protein synthesis by introducing a 10-minute pulse (Figure 2). The pulse stimulation primes PKC-mRNA and stimulates the new synthesis of PKC enzyme. Clearly, a complete model of protein synthesis should account for all the complex molecular processes involved in new protein synthesis, including the availability and assembly of translational machinery. However, this is beyond the scope of this study. Similarly, this model assumes that stable PKC-mRNA is available in the cytosol for new protein synthesis during simulations. Here, we are not attempting to model the transcription of PKC. Our analysis of the PKC life cycle accounts for post-transcriptional events involved in the signaling lifetime of this enzyme. Additionally, we modeled the generation of second-messenger DAG through a brief pulse. This is a simple description of complex processes involved in the generation of second-messenger DAG at the plasma membrane. We are not accounting for all upstream processes such as PIP_2_ and PLC activation prior to the DAG generation at the plasma membrane. The main goal of this study is to analyze the enzymological characteristics of PKC during various phases of its life cycle, not the detailed mechanisms of second-messenger generation and metabolism.

Recent experimental results indicate that PKC adopts an open conformation following second messenger-mediated binding and activation. This state of PKC enzyme is highly sensitive to phosphatases and quickly undergoes PHLPP_1_ and PHLPP_2_ mediated dephosphorylation. The dephosphorylated molecule has reduced thermal stability and, if maintained in this state for long, could be targeted for down-regulation and degradation. Observations also show that hsp70 binds this dephosphorylated species via the turn motif and prevents its association with degradation and down-regulation machinery. Here, our study also analyzes the role of hsp70 during the PKC life cycle especially in rescuing the enzyme from degradation pathways and preventing the down-regulation of active but de-phosphorylated state of the enzyme (Figure 6). Our results show that for endogenous hsp70, a high intensity pulse stimulation of second messenger can lead to complete down-regulation of PKC enzyme (Figure 6 a, b, c & d**: No HSP70 overexpression case**). This occurs because high intensity stimulation generates a larger pool of active and dephosphorylated PKC, thus providing enough time for degradation pathways to interact with PKC and target it for down-regulation (Figure 6d**: No HSP-60 overexpression case**). Our model predicts that if the expression levels of HSP-60/70 are controlled (Figure 6a), it produces a dose-dependent reversal of PKC down-regulation even after the application of a high intensity second messenger stimulation pulse (Figure 6a). We believe this occurs because HSP-60/70 binds with the dephosphorylated PKC molecule and modulates its re-phosphorylation, thus replenishing the enzyme pool in the cytosol. Our results show that heat shock protein overexpression reduces the levels and duration of persistence of the PKC^A^ species (Figure 6d), thus completely or partially restoring the cytosolic enzyme pool.

Interestingly, at first glance the life cycle model of this work suggests the stabilizing roles of PDK1 and HSP70 are functionally equivalent, as both these enzymes are acting on unstable and degradation prone species of PKC isoforms. However, newly synthesized species and dephosphorylated active but mature species are conformationally inequivalent. This model is designed to account for this critical difference, as PDK1 can only bind nascent enzyme. HSP70 is set to bind to mature, active and dephosphorylated form PKC. This model clearly defines the roles of PDK_1_ and HSP70 during the different phases of the PKC life cycle.

Recent experimental results using a PHLPP knock-down indicate that these novel phosphatase isoforms can act as negative regulators of PKC signaling. Data shows that a PHLPP knockdown in colon cancer cell lines is linked to a 3-fold increase in the expression of PKCβII. Similar trends are also observed in breast epithelial cells. These results suggest that cellular levels of PKC can be modulated by PHLPP isoforms. Our model predicts that blocking PHLPP could reverse the second messenger-induced down-regulation of PKC. Our results also predict that degree of PKC down-regulation could be reversed by inhibiting the PHLPP isoforms in a dose-dependent manner (Figure 7 a, b, c, d). This occurs because inhibiting the dephosphorylation pathways will reduce the levels of PKC^A^, thus reducing the propensity of down-regulation (Figure 7d). Our results suggest that PHLPP molecules play a critical role in modulating the signaling lifetime of PKC. Targeting the expression or activity of PHLPP’s could be a key factor in fine-tuning the signaling characteristics and functionality of this family of enzymes.

Some of the assumptions we made to construct this model might still be disputed. The enzymological characteristics of cPKCs are critically regulated by their interactions with scaffold proteins. Previous studies indicate that PKCα interacts with scaffold proteins like PICK_1_, PSD95 and SAP97 [36–40]. It seems that interactions with these proteins could be important to the regulation of structure, function and pharmacological responsiveness of PKC. In the proposed PKC life cycle model, we are not accounting for these interactions. It is possible that, by incorporating these additional interactions, one could further extract more interesting features of the PKC life cycle. These factors could be PKC translocation, anchoring and localization at specific membrane locations. However, it is unlikely that the overall structure and conclusions drawn from this model would be significantly altered. At best, our model is a simplified representation of all the molecular events that occur during the PKC life cycle. Though some kinetic and degradation rates are estimated from previous experimental data, many still are unknown. Therefore, though we are making certain quantitative predictions, further work is needed to precisely estimate all the kinetic rate constants. In parallel, we are also developing a two-compartment model of the PKC life cycle through incorporating membrane translocation and remigration back to cytosol. This version will provide a more realistic representation of the PKC life cycle.

Recent experimental observations suggest that protective action of endogenous heat shock proteins during the PKC lifecycle could be linked to a feed forward regulation [41]. These data sets show that phorbol esters, which stimulate the translocation and activation of cPKC family members, may also enhance the responsiveness and expression of heat shock proteins. This raises an interesting possibility that hsp70 could robustly prolong the lifetime of PKC [41]. Although activation of PKC targets it for dephosphorylation and down-regulation, the same stimuli may also enhance the capacity of heat shock proteins to rescue PKC from degradation pathways. This could be interesting possibility for modulating the gain-of-function (GOF) and LOF in specific pathological conditions however, in the current study we are not incorporating this feed forward protective cascade in our life cycle model. In our proposed life cycle model, the heat shock protein is considered at a fixed expression and activity levels. In our simulations heat shock protein expression is set at a certain value in the initial conditions and remains constant throughout the simulation. In our upcoming version of this model we are planning to incorporate a feed forward protective response in a detailed two-compartment system.

The current model is motivated by experimental observations. However, its assumptions as well as its consequences need to be further tested. One of the key assumptions in our proposed model is that only PHLPP_1_ and PHLPP_2_ are responsible for complete dephosphorylation of active PKC. This assumption is supported by observations that PHLPP_1_ and PHLPP_2_ are linked to dephosphorylation of active PKC. However, it is possible that there are other phosphatases present which may also be contributing to enzyme dephosphorylation. Our model only considers PHLPP1 as the relevant phosphatase in the dephosphorylation of active PKC species. Another assumption we made in regards to PKC degradation only takes place through dephosphorylated active species. However, some observations show that active and phosphorylated species could also degrade directly at the membrane. Here, we are not incorporating this into our model for the sake of simplicity.

## Materials & Methods

### 4.1 Biochemical Reactions

The biochemical reactions that comprise the PKC life cycle (Fig. 1) are based on standard mass action kinetics. The following set of reactions describes the molecular events that take place during the PKC life cycle. PKC is one of the dynamical variables and represents the naïve form of protein kinase C. PKC-mRNA represents the RNA transcript that codes for the PKC protein. PKC-P_A_ represents the PKC protein that is phosphorylated at an activation site during a PDK_1_ mediated event. C.PKC-P_A_ represents a complex of mTROC_2_ and PKC-P_A;_ whereas the complex C.PKC-P_A_.P_H_.P_T_ represents a PKC complex in which PKC is phosphorylated at all three sites: the activation site, turn site, and the hydrophobic motif site. C.PKC-P_A_.P_H_.P_T_ also represents a form of the enzyme that is inactive, but catalytically competent. This form of the enzyme is stored in the cytosol. The PKC.P_A_.P_H_.P_T_ ^A^ form of enzyme represents an active, phosphorylated molecule found at the cell membrane. The dephosphorylated, but active species is represented by PKC^A^. The phosphatase, P, is approximated as a fixed parameter rather than a dynamical variable in order to simplify the simulations. This approximation did not significantly alter the results. PDK_1_ represents the concentration of constitutively active PDK_1_ and is a fixed parameter. Here, ‘T’ represents the concentration of polyribosome and is a fixed parameter. The mTORC_2_, Ca^+2^/DAG, PHLPP & HSP70 are all fixed parameters and represent the concentrations of complex mTORC2, secondary-messenger DAG, phosphatases PHLPP1 & PHLPP2, and the concentration of heat shock protein 70, respectively.

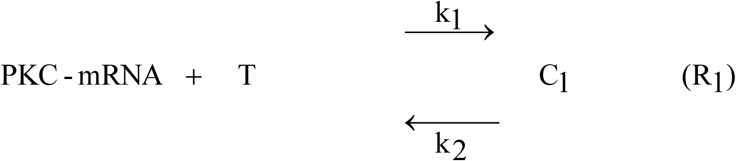

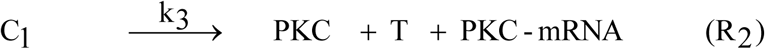

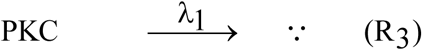

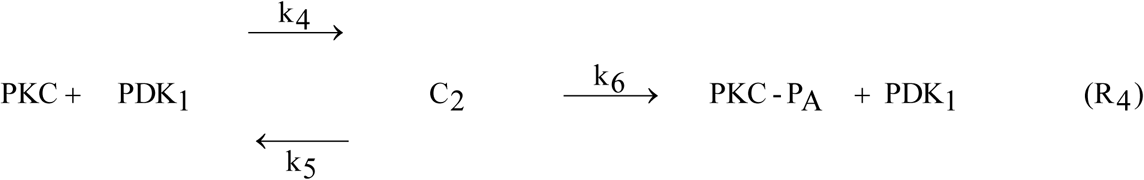

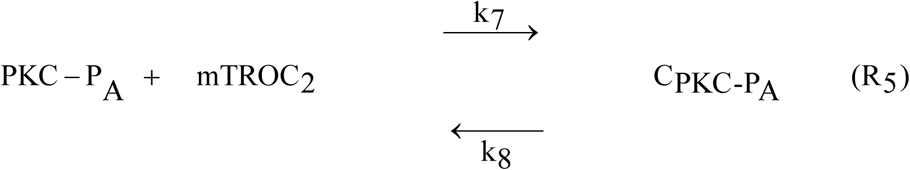

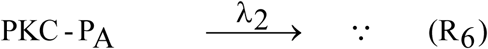

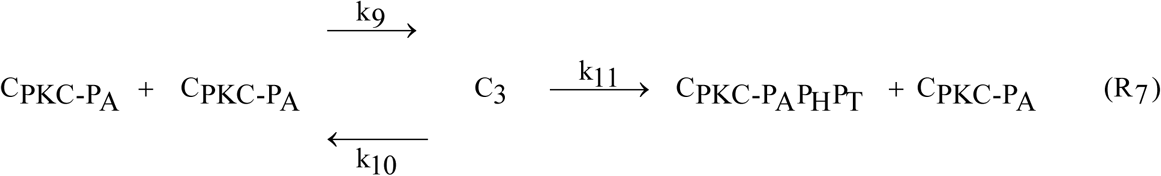

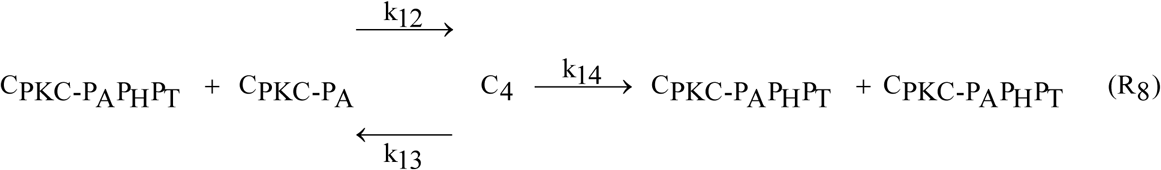

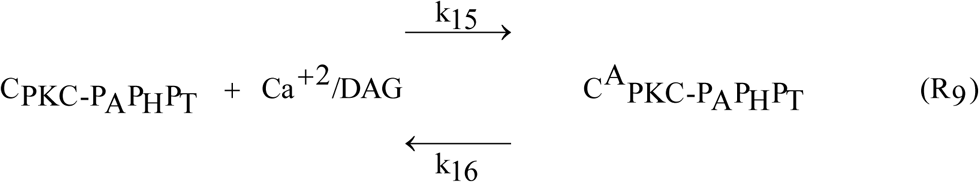

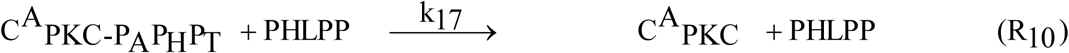

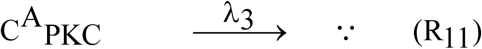

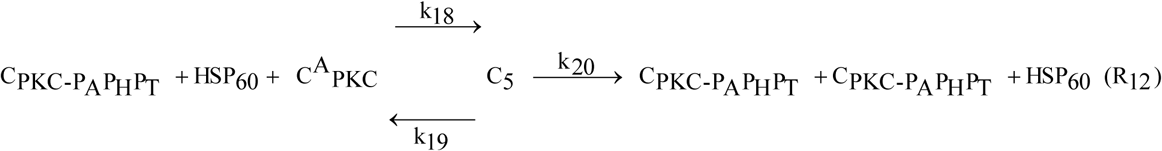

The PKC life cycle model described by the biochemical reaction equations (Figure 1, R_1_-R_12_) can be perturbed either through stimulation of protein synthesis or through secondary-messenger Ca^+2^/DAG-modulated activation. New PKC synthesis is described by equations R1 and R2. (Figure 1). These reactions show that when new protein synthesis is stimulated, the PKC transcript is loaded with polyribosome “T,” as described by R_1_ (Figure 1). The loaded transcript “C_1_” can then undergo translation to generate additional PKC protein (Figure 1). During induction, the signals “k_3_” from multiple upstream kinases act on this loaded transcript and initiate the *de-novo* synthesis of naïve PKC from its transcript (Figure 1). This translation event is described by the biochemical reaction event R_2_ (Figure 1). The naïve PKC molecule is highly unstable and can quickly degrade via degradation pathways, as described by the biochemical reaction event R_3_ (Figure 1). However, the naïve, unstable PKC molecule has a strong affinity for constitutively active kinases (i.e., PDK_1_), and undergoes maximum phosphorylation at its activation site (i.e., P_A_) as a result (Figure 1, reaction event R_4_). The phosphorylated enzyme (PKC.P_A_) forms a biochemical complex (C.PKC.P_A_) with the mTORC_2_ complex (Figure 1, reaction event R_5_). Phosphorylation at activation site P_A_ reduces the degradation rate of naïve enzyme because PKC.P_A_ is more stable and degrades more slowly as described by R_6_. In contrast, λ_2_ represents the degradation rate of this enzyme species (Figure 1). The C.PKC.P_A_ complex stimulates autophosphorylation at the hydrophobic sites of the PKC enzyme. This process generates the fully mature, auto-inhibited, catalytically competent species C.PKC.P_A_.P_H_.P_T_. These autophosphorylation events are described through equations R_7_ & R_8_ (Figure 1). The inactive, auto-inhibited, catalytically competent species is stored at different locations in the cytosol. Following membrane receptor stimulation, second messengers prompt this species to migrate from the cytosol to the membrane, where a second messenger activates it (Figure 1, reaction event R_9_). The active C.PKC.P_A_.P_H_.P_T_ ^A^ form of the enzyme is prone to rapid dephosphorylation by phosphatases PHLPP_1_ and PHLPP_2_. These dephosphorylation events are described by equation R_10_ (Figure 1). Upon dephosphorylation, the active form of the enzyme, C.PKC^A^, is prone to degradation, and is thus targeted by degradation machinery for removal (Figure 1, reaction event R_11_). Heat shock proteins HSP-60 and HSP-70 act on the active and dephosphorylated form of the enzyme, rescuing and recovering PKC from degradation pathways and restoring the pool of inactive, catalytically competent PKC in the cytosol (Figure 1, reaction event R_12_).

### 4.2 Induction

During simulations, the induction of new proteins was simulated by a 10-minute pulse of protein synthesis, which produced an increase in the total PKC protein concentration.

### 4.3 Temporal Dynamics

The differential equations resulting from above interactions were integrated through nonlinear solvers using MATLAB (MathWorks). The dynamical coefficients’ values were estimated from limited experimental data. Unknown rate constants were scaled to obtain dynamics that were comparable to experimental values [1–9,18–21,23–24]. Unless otherwise stated, all of the molecular concentrations in the model are expressed as ng/ml and time is represented in minutes.

## Acknowledgments

Special thanks to Miss Kaneez Fatima for support

## Supplementary Material 1: Supplementary Figures

**Figure S1:**
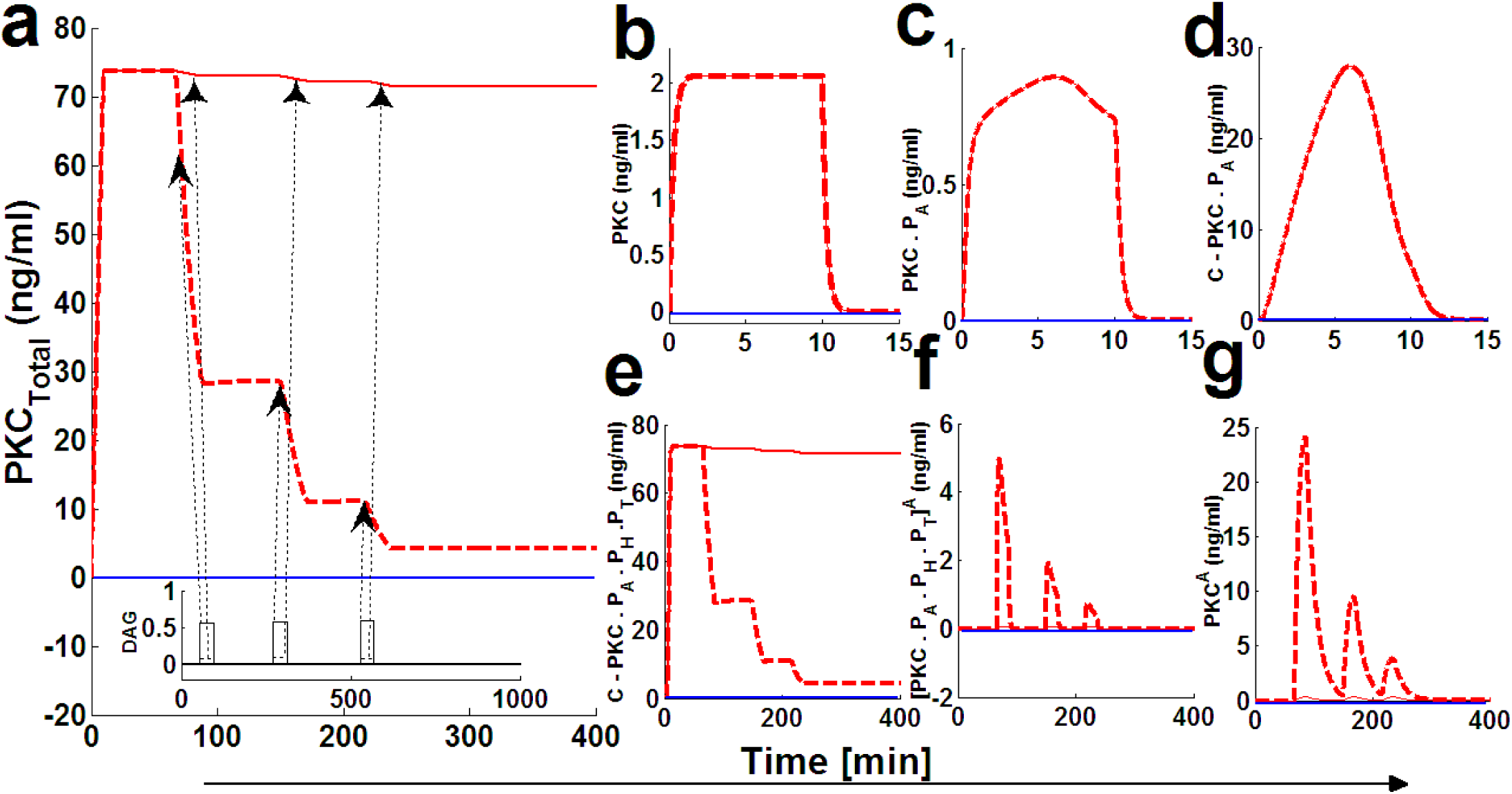
The numerical simulations comparing the down-regulation characteristics of cPKC during sequential application of three lower intensity and higher intensity second messenger pulse stimulations. These results indicate that initial application of a 10-minute protein synthesis pulse leads to the generation of a naïve, and unstable enzyme which is quickly stabilized through constitutive phosphorylation events at the activation, turn and hydrophobic sites. Once stable and phosphatase/proteasome resistant this inactive form of cPKC is stored in cytosol. Here, dashed red lines indicate stimulation (protein synthesis as well as second-messenger) and blue solid lines indicate non-stimulated condition. (a) Total PKC enzyme during protein synthesis and second messenger mediated stimulation. These results show that application of a brief pulse (**dashed-line region**) shows the quick generation of cPKC enzyme and stabilization. These results also show that at later time points (i.e., 50, 150 and 200 minutes in the stimulation) the application of three lower intensity and three higher intensity second messenger pulse (15-minute pulse mimicking Ca^+2^/DAG: lower intensity DAG is set at 0.0005 nM and higher intensity DAG is set at 0.05nM) leads to the down-regulation of cPKC. However, the degree of down-regulation in case of lower intensity stimulation is minimal in contrast, degree of down-regulation is quite significant in case of higher intensity stimulation. (b) concentration of naïve, newly synthesized unstable form of cPKC. (c) concentration of PKC species phosphorylated at activation site. (d) concentration of PKC complex C.PKCP_A_. (e) concentration of PKC complex, phosphorylated at all three sites i.e., activation, turn and hydrophobic sites. (f) concentration of active complex PKC.P_A_.P_H_.P_T_.^A^ after second messenger binding. Again, these results indicate that activation levels of this complex are dependent on the levels of second-messenger stimulation. (g) concentration of dephosphorylated but active PKC^A^ molecule.

### Supplementary Material 2

**Table 1.**
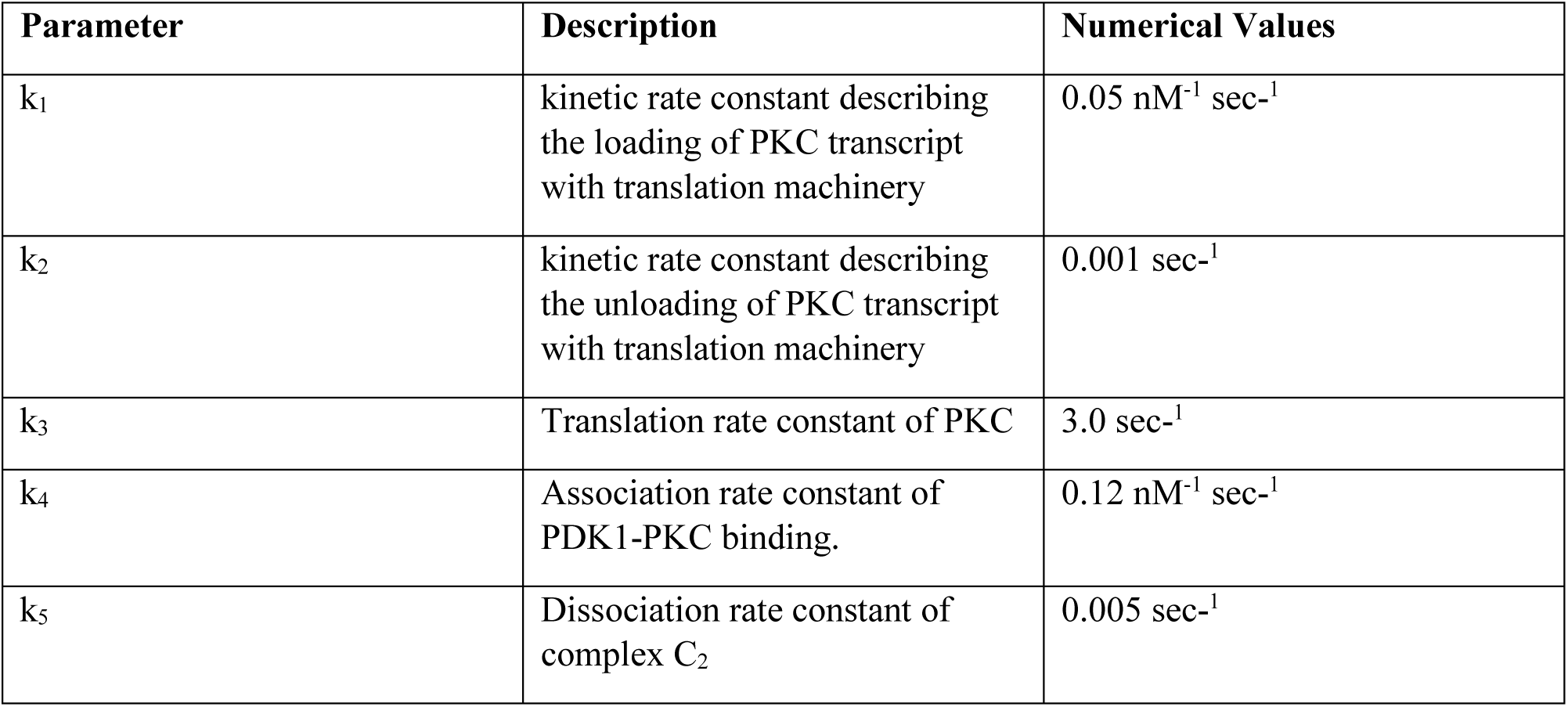
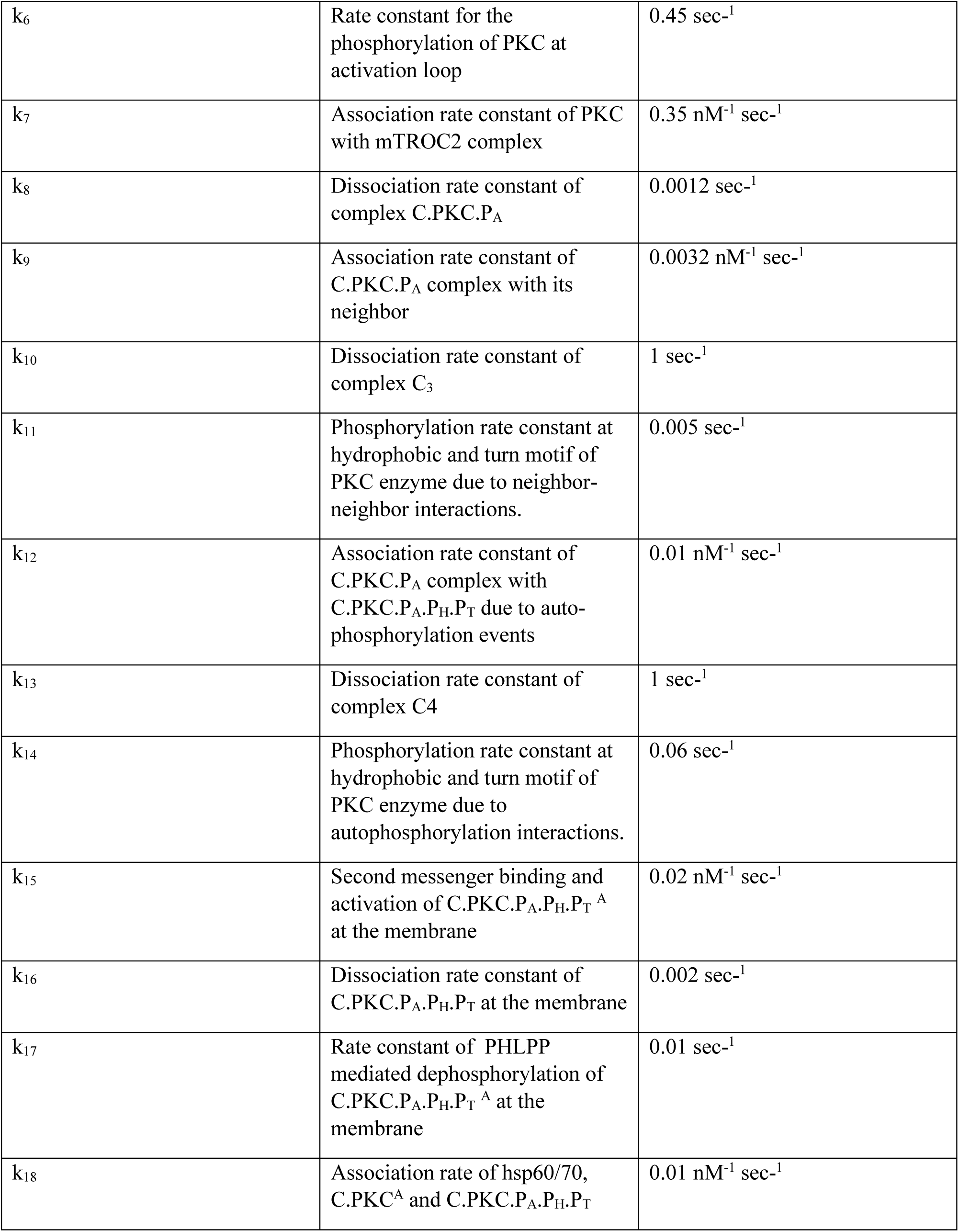
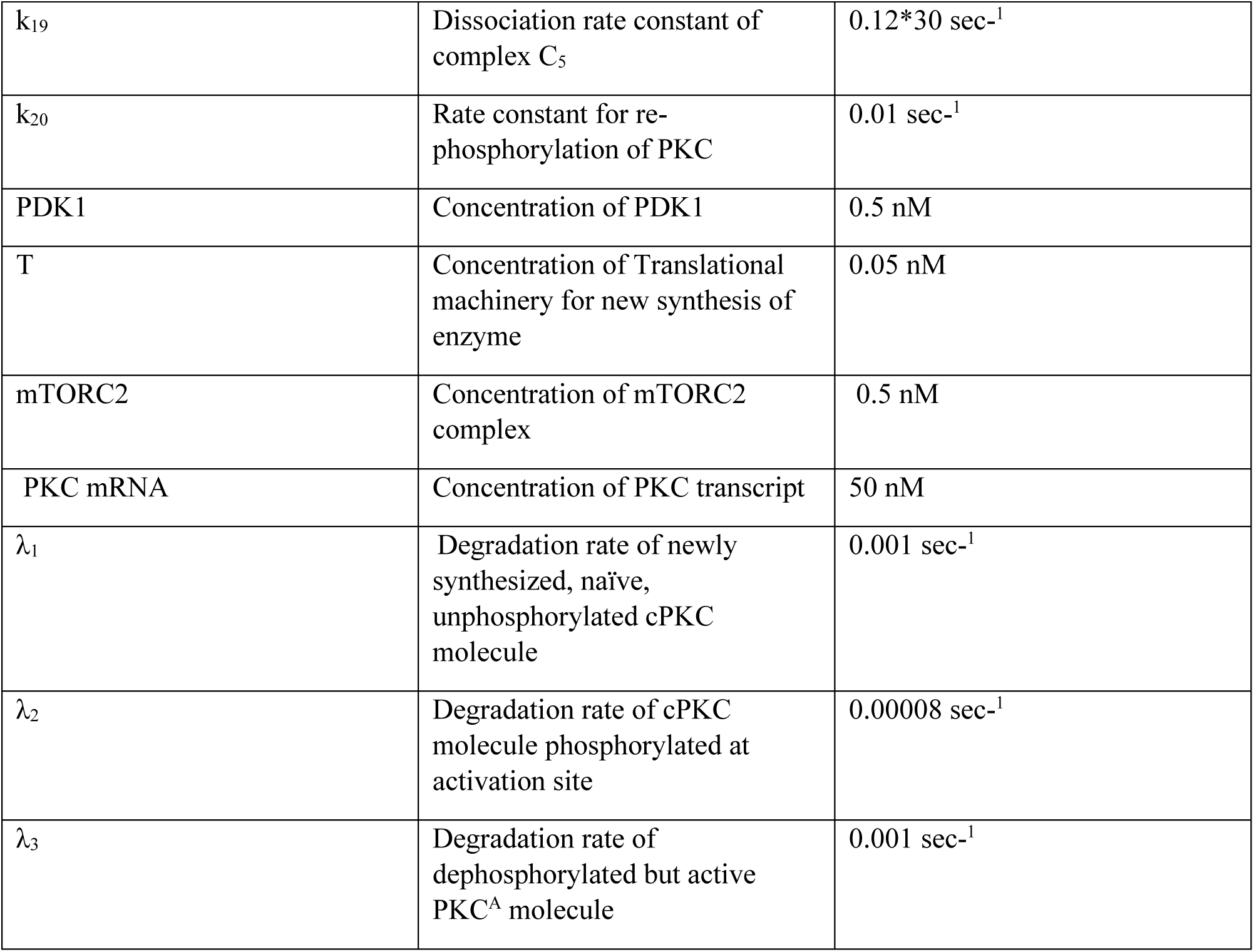
Numerical values of biochemical rate parameters representing the molecular model describing the PKC life cycle

| Parameter | Description | Numerical Values |
| --- | --- | --- |
| $k_1$ | kinetic rate constant describing the loading of PKC transcript with translation machinery | $0.05 \text{ nM}^{-1} \text{ sec}^{-1}$ |
| $k_2$ | kinetic rate constant describing the unloading of PKC transcript with translation machinery | $0.001 \text{ sec}^{-1}$ |
| $k_3$ | Translation rate constant of PKC | $3.0 \text{ sec}^{-1}$ |
| $k_4$ | Association rate constant of PDK1-PKC binding. | $0.12 \text{ nM}^{-1} \text{ sec}^{-1}$ |
| $k_5$ | Dissociation rate constant of complex $C_2$ | $0.005 \text{ sec}^{-1}$ |
| $k_6$ | Rate constant for the phosphorylation of PKC at activation loop | $0.45 \text{ sec}^{-1}$ |
| $k_7$ | Association rate constant of PKC with mTROC2 complex | $0.35 \text{ nM}^{-1} \text{ sec}^{-1}$ |
| $k_8$ | Dissociation rate constant of complex C.PKC.P<sub>A</sub> | $0.0012 \text{ sec}^{-1}$ |
| $k_9$ | Association rate constant of C.PKC.P<sub>A</sub> complex with its neighbor | $0.0032 \text{ nM}^{-1} \text{ sec}^{-1}$ |
| $k_{10}$ | Dissociation rate constant of complex C<sub>3</sub> | $1 \text{ sec}^{-1}$ |
| $k_{11}$ | Phosphorylation rate constant at hydrophobic and turn motif of PKC enzyme due to neighbor-neighbor interactions. | $0.005 \text{ sec}^{-1}$ |
| $k_{12}$ | Association rate constant of C.PKC.P<sub>A</sub> complex with C.PKC.P<sub>A</sub>.P<sub>H</sub>.P<sub>T</sub> due to auto-phosphorylation events | $0.01 \text{ nM}^{-1} \text{ sec}^{-1}$ |
| $k_{13}$ | Dissociation rate constant of complex C4 | $1 \text{ sec}^{-1}$ |
| $k_{14}$ | Phosphorylation rate constant at hydrophobic and turn motif of PKC enzyme due to autophosphorylation interactions. | $0.06 \text{ sec}^{-1}$ |
| $k_{15}$ | Second messenger binding and activation of C.PKC.P<sub>A</sub>.P<sub>H</sub>.P<sub>T</sub><sup>A</sup> at the membrane | $0.02 \text{ nM}^{-1} \text{ sec}^{-1}$ |
| $k_{16}$ | Dissociation rate constant of C.PKC.P<sub>A</sub>.P<sub>H</sub>.P<sub>T</sub> at the membrane | $0.002 \text{ sec}^{-1}$ |
| $k_{17}$ | Rate constant of PHLPP mediated dephosphorylation of C.PKC.P<sub>A</sub>.P<sub>H</sub>.P<sub>T</sub><sup>A</sup> at the membrane | $0.01 \text{ sec}^{-1}$ |
| $k_{18}$ | Association rate of hsp60/70, C.PKC<sup>A</sup> and C.PKC.P<sub>A</sub>.P<sub>H</sub>.P<sub>T</sub> | $0.01 \text{ nM}^{-1} \text{ sec}^{-1}$ |
| $k_{19}$ | Dissociation rate constant of complex $C_5$ | $0.12 \cdot 30 \text{ sec}^{-1}$ |
| $k_{20}$ | Rate constant for re-phosphorylation of PKC | $0.01 \text{ sec}^{-1}$ |
| PDK1 | Concentration of PDK1 | 0.5 nM |
| T | Concentration of Translational machinery for new synthesis of enzyme | 0.05 nM |
| mTORC2 | Concentration of mTORC2 complex | 0.5 nM |
| PKC mRNA | Concentration of PKC transcript | 50 nM |
| $\lambda_1$ | Degradation rate of newly synthesized, naïve, unphosphorylated cPKC molecule | $0.001 \text{ sec}^{-1}$ |
| $\lambda_2$ | Degradation rate of cPKC molecule phosphorylated at activation site | $0.00008 \text{ sec}^{-1}$ |
| $\lambda_3$ | Degradation rate of dephosphorylated but active PKC<sup>A</sup> molecule | $0.001 \text{ sec}^{-1}$ |

